# Temporal predictions as motor readouts of sensory predictions

**DOI:** 10.1101/2025.06.18.660330

**Authors:** Aaron Kaltenmaier, Quirin Gehmacher, Peter Kok, Matthew H. Davis, Clare Press

## Abstract

When will I see something and what will it be? Temporal predictions are crucial for adaptive interaction with the environment and are typically accompanied by predictions about sensory content, yet these two types of ‘what’ and ‘when’ predictions are usually studied separately. Specifically, oscillatory phase-coupling (or ‘entrainment’) has been proposed to align our neural sensitivity with likely moments of stimulus appearance, however these accounts ignore that predictions about when something will appear are usually accompanied by predictions about what it will be. Thus, temporal predictions may not enhance all sensory processing but rather modulate particular channels encoding predicted content. We here demonstrate oscillatory phase-coupling in vision and show how it relates to content-specific encoding. In a magnetoencephalography (MEG) study, participants observed rhythmic Gabors at 1.33 or 2 Hz with predictable orientations. They judged the timing or orientation of a delayed probe which manipulated the requirement to covertly maintain the sequence rhythm. We found sustained oscillatory phase-coupling to the entrained rhythm in motor areas specifically when participants judged stimulus timing, where its extent was associated with perceptual performance. Meanwhile, neural decoding revealed content predictions in early visual areas (‘what’) that fluctuated in line with temporal predictions (‘when’). These temporally-specific content predictions appeared regardless of task instruction but were correlated and phase-aligned with the motor phase-coupling during timing judgements. These findings suggest that temporal predictions may be derived from motor readouts of temporally-specific sensory predictions, with broad implications for our understanding of entrainment and prediction, and how we represent time more generally.

## Results

Thirty human participants observed rhythmic sequences of 7-8 flashing Gabor patches that were predictable in both time and content. Specifically, they were presented at 1.33 or 2Hz and rotated with a fixed angle and direction. These sequences were followed by a stimulus-empty interval and finally a probe (Figure 1a). We examined temporal predictions generated by the visual rhythms in the empty interval between sequence and probe to ensure that effects are uncontaminated by stimulus presentation (see below; (1)). We manipulated the necessity to maintain temporal information during the empty interval by contrasting a ‘timing’ with an ‘orientation’ task. In the timing task, participants judged whether the delayed probe appeared at the timepoint predicted from a continuation of the preceding sequence (i.e. on the beat of the preceding rhythm). In a separate block of orientation task trials, participants instead judged whether the delayed probe had an orientation that would have immediately followed the final orientation of the sequence (i.e. continuing the sequence as if uninterrupted). Probes were either as expected or could appear half a frequency cycle too early or too late and be oriented either one step further or less than expected. Thus, we could present identical stimuli across the two tasks but simply alter which feature was judged.

**Figure 1.**
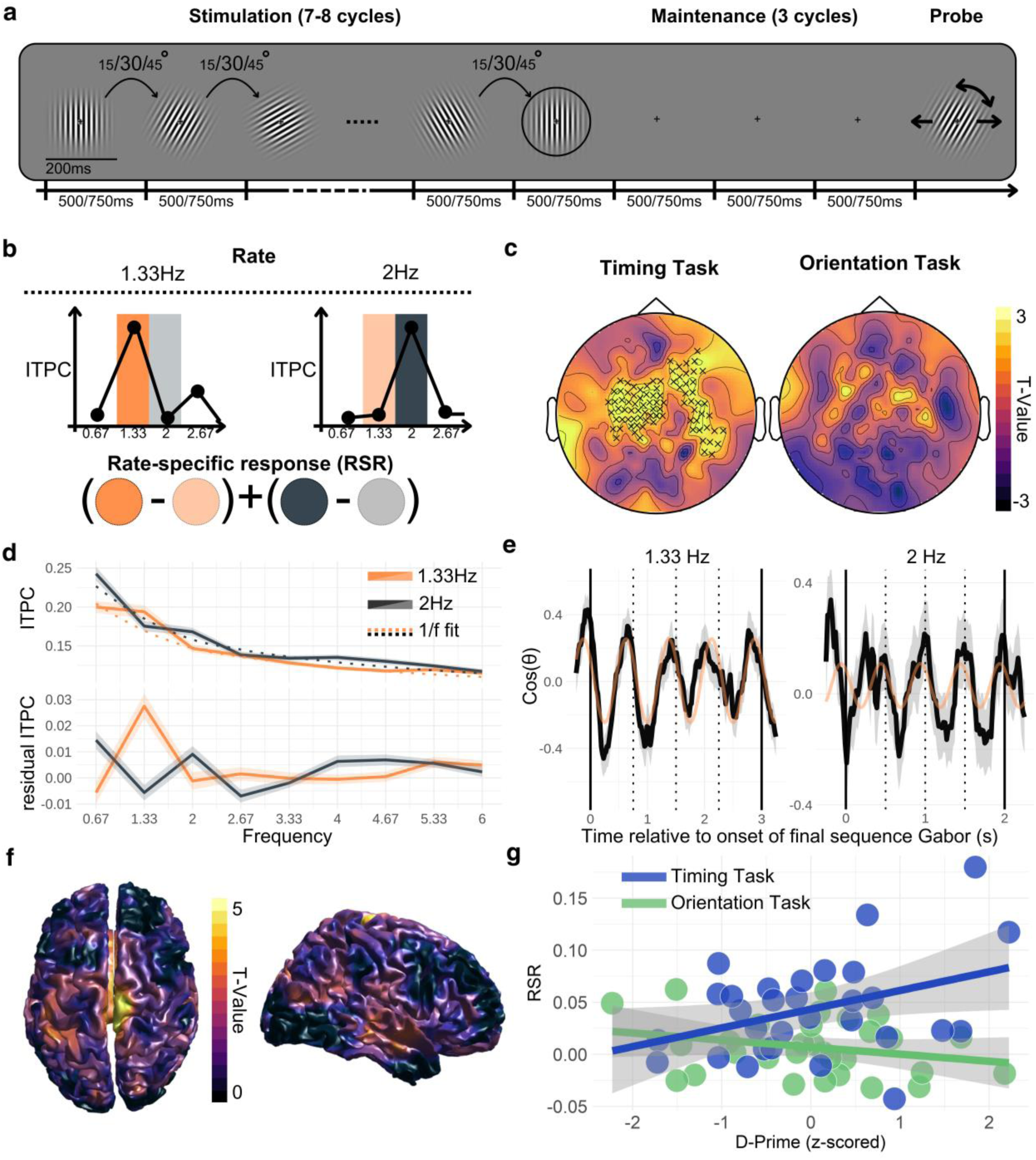
Design and oscillatory rate specific response (RSR) analysis findings. **a** Participants observed rhythmic (1.33 or 2Hz) sequences of rotating Gabor patches before judging a delayed probe’s orientation or timing. **b** RSR was determined by contrasting intertrial phase consistencies (ITPCs) of stimulated and non-stimulated rates. **c** Permutation test t-value topographies for the RSR during the Maintenance window, separately for the two tasks. Sensors marked with a cross belonged to the significant cluster found for a given condition. **d** ITPC by frequency in the timing task, separated by stimulation rate and averaged over sensors belonging to significant cluster in c. Top and bottom panels display ITPC without and with removal of 1/f activity (dotted line in top panel, see Methods). Solid line represents group mean, shaded region the standard error of the mean. **e** Cosine-transformed phase in the timing task for the rate-congruent frequency (Left: 1.33Hz, Right: 2Hz), averaged over sensors belonging to cluster in c. Solid trend line represents group mean, shaded region the standard error of the mean. Orange lines show best-fit cosine of the group mean. Solid vertical lines represent the onset of the final sequence Gabor and probe respectively. Dotted vertical lines represent the rate-specific predicted onsets during the stimulus-empty window. **f** Same as **c**’s timing task but source-localised to motor cortex using an LCMV beamformer. **g** Correlations between rate-specific neural activity averaged across the sensors belonging to the significant cluster in **c** and task performance, separately for the orientation and timing task. Each circle represents a single participant (N = 30) in the given task.

Participants were able to perform both tasks at both stimulus presentation rates with above-chance sensitivity (one-sample t-test of d-primes against 0: all p<0.001). In the orientation task, average d-primes in 1.33Hz and 2Hz blocks were 1.516 (SEM = 0.126) and 1.348 (SEM = 0.102) respectively. Performance in 1.33Hz and 2Hz timing task blocks was 0.667 (SEM = 0.111) and 0.425 (SEM = 0.099) respectively. A 2 (Rate: 1.33Hz, 2Hz) by 2 (Task: Orientation, Timing) repeated-measures ANOVA confirmed main effects of both rate (F(1,29) = 10.611, p = 0.003) and task (F(1,29) = 50.299, p < 0.001), without an interaction between the two (F(1,29) = 0.329, p = 0.571; see Supplementary Information for demonstration that task difficulty differences could not account for the neural effect differences that follow; Table S1, Figure S1).

### Visual temporal predictions are encoded in the oscillatory phase of motor regions

In line with previous auditory work, we probed for oscillatory phase-coupling by contrasting inter-trial phase consistency (ITPC) between the two rates of stimulus presentation. We computed a rate-specific response (RSR) at each MEG sensor that precisely captures how much stronger the ITPC is at the trial’s stimulated compared to non-stimulated frequency (1) (Figure 1b). To distinguish oscillatory phase-coupling from evoked responses during rhythmic stimulation (2–4), we tested for an RSR in a stimulus-empty window following rhythmic sequences and before the probe (see Supplemental Information, Figure S2, for the RSR during stimulation).

We found a cluster of significant RSR in the timing task that had a central topography spanning frontal, parietal and temporal sensors bilaterally (cluster-corrected p < 0.001; summed t = 239.426; 86 sensors; Figure 1c). However, no significant RSR cluster was found during the orientation task blocks (Figure 1c). Comparing the two tasks directly, a central cluster emerged in which maintained RSR values were significantly higher in the timing than in the orientation task (cluster-corrected p = 0.019; summed t = 44.93; 17 sensors). All sensors belonging to this task difference cluster were also part of the cluster found in the timing task alone. Using an LCMV beamformer in combination with a probabilistic atlas ((5), see Methods), we localized RSR during the timing task to the right paracentral lobule, specifically the supplementary motor area, which has recently been implicated in the preparation, imagination and suppression of movement ((6–8), Figure 1f; note that we were unable to find neural evidence of overt movement that may explain this response (Figure S3)).

Examining each stimulation rate’s individual contributions to this cluster, we separated the timing task RSR into its separate 1.33Hz and 2Hz components that are summed over in the calculation of RSR (Figure 1b). We found stimulation-specific ITPC in both the 1.33Hz (M = 0.033, t(29) = 6.227, p < 0.001) and 2Hz (M = 0.010, t(29) = 2.408, p = 0.023) condition that was significantly larger in the former than in the latter (M = 0.023, t(29) = 5.119, p < 0.001). As in the RSR, stimulation-independent spectral differences are controlled for in these components as they contrast phase consistency during rate-congruent and rate-incongruent visual stimulation. In line with past research (9), this difference between two stimulation rates may aid to explain the better behavioural timing task performance in the 1.33 compared to the 2Hz condition. Examining raw phase over time additionally revealed that activity indeed oscillated in phase with temporal predictions (Figure 1e, also see S4).

To establish whether the RSR is associated with perception, we next extracted participant-specific RSR values from the timing task cluster and related these to behavioural performance (d-prime). We averaged over sensors to create a single value per participant, separately for each task, and found a significant correlation between RSR responses and timing task performance (r = 0.385, p = 0.036), while no such correlation over participants was found for the orientation task (r = -0.251, p = 0.182; Figure 1g) – in line with the absence of an RSR in this condition. Comparing the two correlation coefficients indeed revealed a large difference between the two (z = 2.460, p = 0.010).

These findings show that temporal predictions were maintained through oscillatory phase-coupling in motor areas, specifically when participants were required to judge the timing of events, and that this oscillatory coupling was associated with the fidelity of timing judgements.

### Content predictions in early visual areas are synchronized to predicted timepoints regardless of task instruction

We next asked about the neural representation of predicted sensory content and the degree of temporal precision in this signal. We again focused on the window between sequence and probe, during which predicted information can be examined independently of presented stimuli. Here, we asked whether the orientations that *would have* been presented had the sequence continued were represented and *at precisely the right moment in time* (Figure 2a). Such an effect would consequently imply that, rather than overall neural amplitude changes as probed in the previous analysis, neural activity that represents predicted sensory content fluctuates in-time with rhythmic temporal predictions.

**Figure 2.**
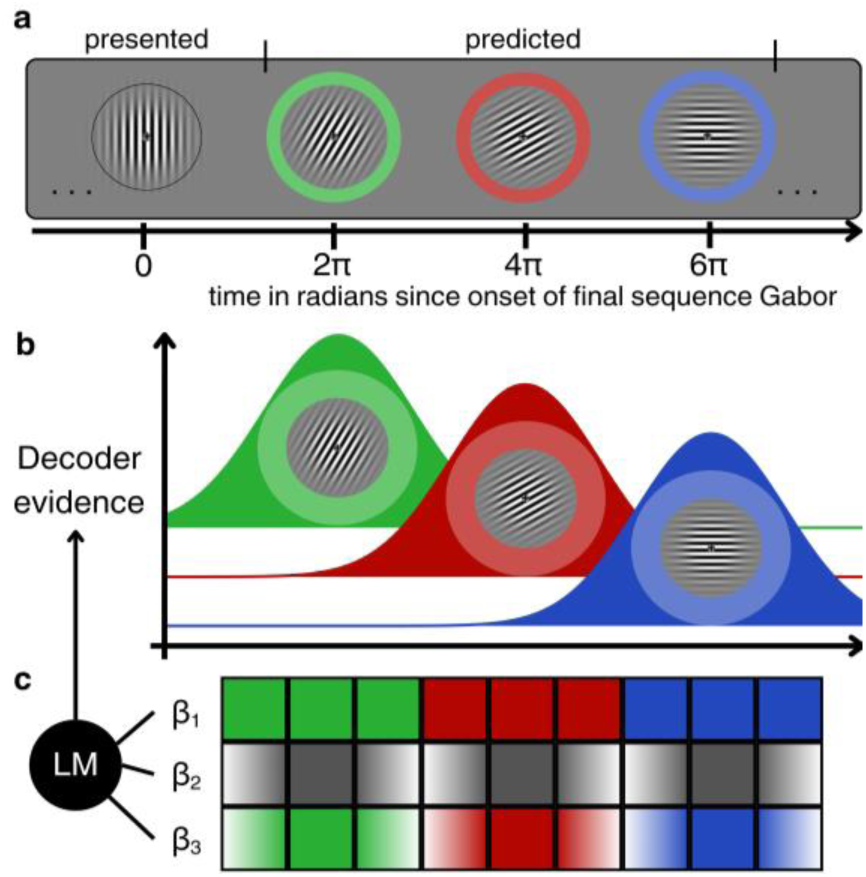
Decoding-based analysis of content-specific temporal predictions. **a** Extrapolating each trial’s rhythmic sequence (see Figure 1a) into the Maintenance window yields one predicted orientation for each of the window’s three ‘empty’ cycles. Relative to the onset of the sequence’s final Gabor, these cycles begin at 2, 4 and 6π rate-specific radians (i.e. if the rate is 2Hz, 2π = 500ms). **b** Neural representation of each of the three predicted orientations should be maximal at their respective predicted timepoint of appearance and therefore temporally-distinct from each other. We use time-resolved decoding (inverted encoding model, IEM) of Gabor orientation to capture these representations. **c** We predicted decoder evidence at each timepoint from content predictions (β_1_), temporal predictions (β_2_) and their interaction (β_3_) using a linear model (LM).

To test this possibility, we trained an inverted encoding model (IEM, see Methods, also (10,11)) to convert MEG sensor-level activity into representations in an orientation feature space. We established the neural origin of the feature-specific signal by source-localising the trained weight matrix of the IEM. In line with past research into predictive content-encoding (10–12), we found contributions originating in occipital cortices and maximal along the left calcarine sulcus (Figure 3a).

**Figure 3.**
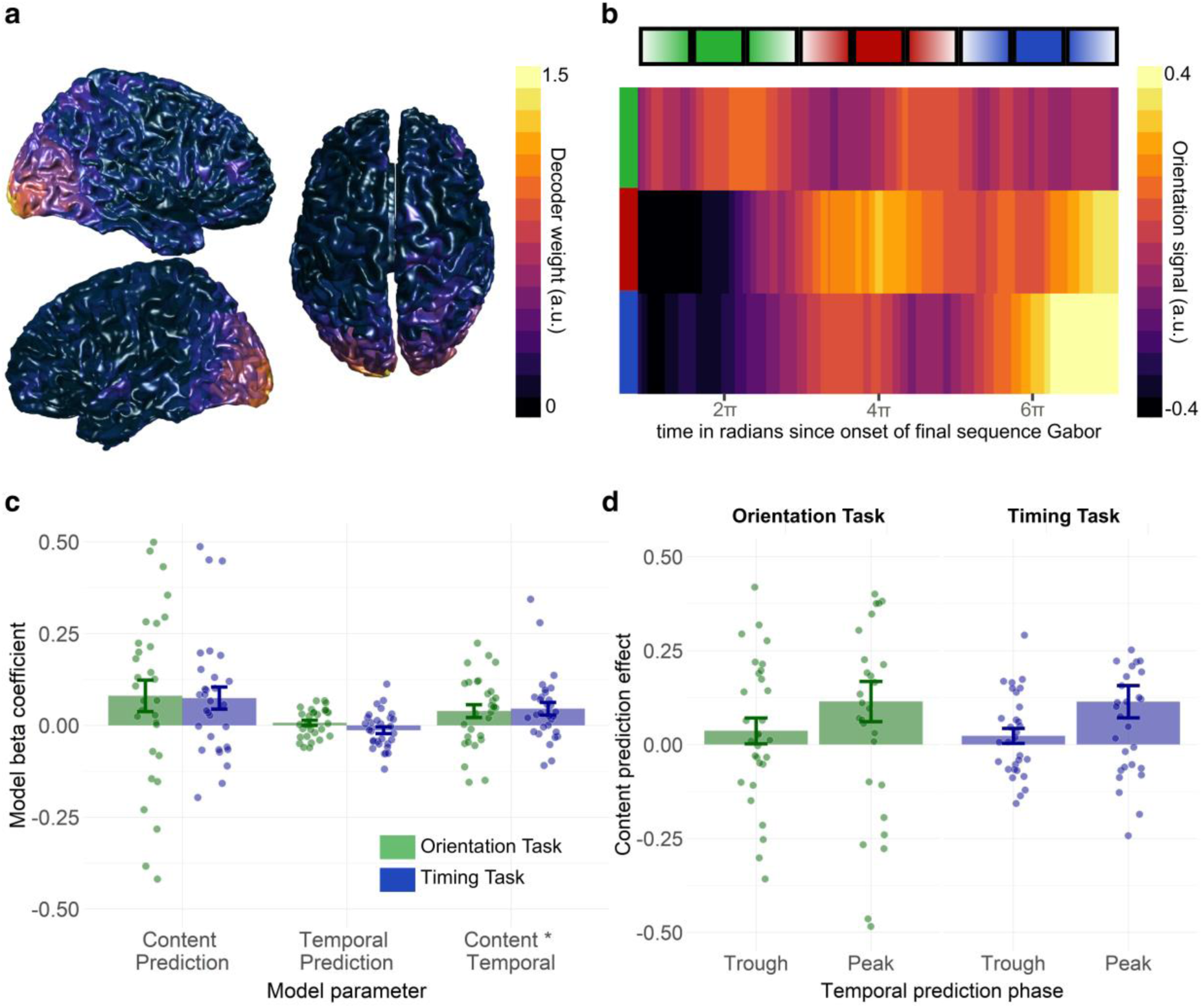
Content-specific temporal predictions. **a** Source-localised inverted encoding model (IEM) weights reveal occipital origin of feature-specific signals. **b** Decoded orientation signal of the three expected orientations (green, red, blue), averaged over all participants and tasks. Horizontal bar atop represents the model interaction term for comparison. **c** Linear mixed effect models were fitted to each participant, separately for each task. Model betas reveal significant group-level effects in both tasks (green: orientation task, blue: timing task) for the content prediction and interaction terms. Error bars reflect the standard error of the group mean. **d** Estimated marginal means illustrate the interaction between content and temporal predictions. The effect of content predictions on decoding evidence (i.e. the difference between neural representation strength of predicted and non-predicted orientations) was significantly larger at the peaks than at the troughs of temporal predictions. Peaks represent the predicted onsets (ticks on x-axis in **b**, cos(t) = 1) and troughs the half-cycles in-between (cos(t) = -1).

We then used the trained model to extract time-resolved decoding evidence for the three orientations predicted in the three rate-specific cycles of each trial’s stimulus-empty window (see above). Using this output and a simple linear mixed effects model (Figure 2b, also see Methods), we tested whether decoding evidence – as a measure of representational strength – changed depending on which orientation was predicted (content predictions) and at what timepoint (temporal predictions). For each of the window’s three rate-specific cycles, one predictor (ß_1_) assessed the neural representation of predicted content, without consideration for *when* within the cycle it was predicted to appear. Note that this first predictor therefore captures a broad, cycle-by-cycle stimulus prediction but does not link it to the precise temporal predictions cued by the preceding rhythm. In contrast, predictor ß_2_ captured whether representation of orientation information in general - regardless of which orientation is predicted – peaked at precisely the moment of stimulus expectation, with sinusoidally graded troughs around it. Combining these two regressors, we also assessed temporally-precise content predictions: the interaction term ß_3_ captured whether the neural representation of predicted content fluctuated in a temporally-precise manner (Figure 2c).

We found that predicted content was represented more strongly than unpredicted content (*b*_1_ = 0.072; *t*(29.026) = 2.399; *p* = 0.023; *d* = 0.45), and that this effect interacted with the phase of the rhythmic temporal prediction (*b*_3_ = 0.042; *t*(29.253) = 2.734; *p* = 0.011; *d* = 0.51; Figure 3b, 3c). Specifically, the difference in representational strength between predicted and unpredicted sensory content was significant at the peaks (*M* = 0.114; *t*(29) = 2.872; *p* = 0.008; *d* = 0.53), but not in the troughs of the rhythmic temporal prediction (*M* = 0.030; *t*(29) = 1.348; *p* = 0.188; *d* = 0.25) (Figure 3d). A main effect of temporal prediction on overall representational strength regardless of content predictions was not found (*b*_2_ = −0.003; *t*(29.162) = −0.451; *p* = 0.655; *d* = 0.08). In stark contrast to the phase-coupling analysis, these effects did not differ between the two tasks (all task interactions p > 0.15), neither did task have a main effect (*b*_4_ = −0.035; *t*(29.007) = −0.490; *p* = 0.628; *d* = 0.09, note that all statistical patterns in these above analyses were replicated if including the three betas in separate, rather than combined, models).

Together, these findings suggest that, in early visual regions, temporal predictions do not produce a uniform increase in neural activity but specifically boost the representation of predicted content at the times when it is predicted to appear (Figure 3d). This content-specific prediction signal thus shows a high level of temporal precision; appearing at the specific moments when visual content is predicted.

### Temporal specificity of content predictions scales with motor phase-coupling

Finally, we asked how the temporal precision of visual content predictions relates to phase coupling in motor regions. For each participant, we quantified the temporal specificity of content predictions as the difference in the content prediction effect (representation of predicted – unpredicted orientations) between the peaks and troughs of temporal prediction (see Figure 3d). Importantly, this temporal specificity was found in both the orientation (M = 0.078, t(29) = 2.205, p = 0.036) and the timing (M = 0.091, t(29) = 2.644, p = 0.013) task (Figure 3d). In the timing task for which we found reliable phase coupling of motor activity to the timing of predicted stimuli, we found a strong relationship between the strength of this phase coupling and the temporal specificity of visual content predictions (r = 0.402, p = 0.027, Figure 4a). Meanwhile, even though we found equally strong temporally-specific content prediction in the orientation task, there was no relationship between this effect and the rate-specific motor effect (*r* = −0.305, *p* = 0.101), perhaps due to the general lack of this signal in the orientation task (Figure 4a). The two tasks’ correlation coefficients again differed from one another (z = 2.725, p = 0.003; note that all statistical patterns in these above analyses were replicated if defining the content prediction effect simply according to predicted orientations, or if using the interaction beta values instead). These results suggest a neural asymmetry whereby temporally-specific visual content predictions appear regardless of whether one is explicitly required to judge time; whereas oscillatory motor responses are only apparent when this is an explicit task requirement (see Figure 1c).

**Figure 4.**
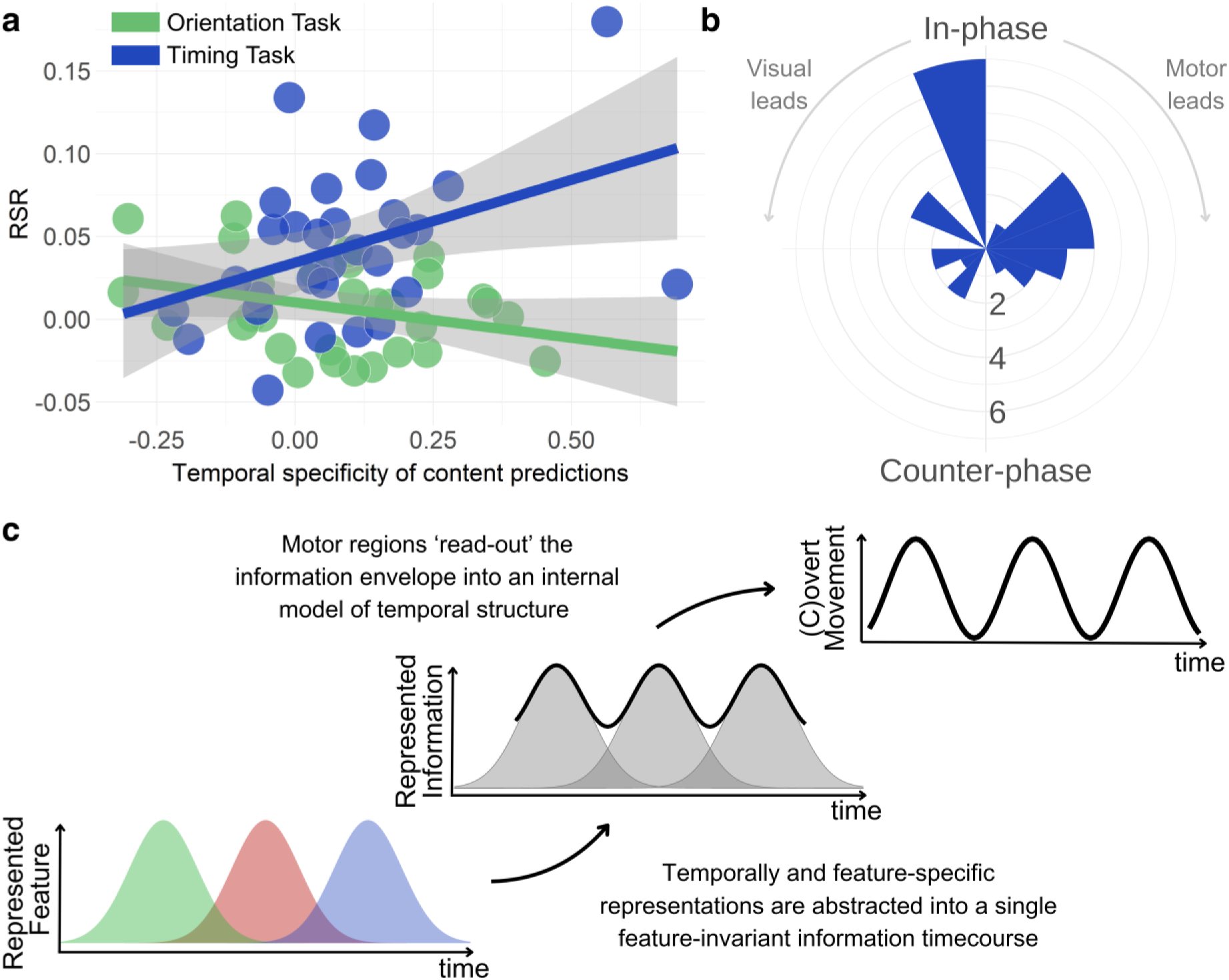
A link between visual content-specific predictions and motor temporal phase-coupling. **a** Correlation between the strength of the visual decoding evidence at peaks vs troughs and the rate-specific response. This relationship can be seen in the timing task, where a group-level RSR was obtained, and not in the orientation task where it was not. **b** Polar histogram of participant-wise lags between content-specific visual activity and rate-specific motor phase in the timing task. In-phase and counter-phase refer to visuomotor lags of 0 and +/-π respectively. **c** Proposed mechanism based on our findings. Activity in sensory processing areas represents predicted stimuli in a temporally and feature-specific manner, which yields an envelope of predicted, information-dense timepoints. Motor regions maintain this envelope and use it to flexibly coordinate temporally-sensitive processing.

Complementing this cross-participant analysis, we also investigated whether the visual and motor signals were temporally aligned within participants in the timing task. This is important since a large motor RSR could theoretically reflect neural activity consistently oscillating in counter-phase to temporal predictions and would thus temporally oppose the neural content prediction effect. We first used a lagged-cosine regression approach to quantify the temporal profiles of the rate-specific motor phase and the content prediction effect in visual areas (see Methods, Figure S4). To estimate the degree of temporal alignment between these two signals, we consequently computed the time lag that yielded the highest cross-correlation between the two neural signatures for each participant. A V-test against the alternative of zero revealed that these visuomotor lags were not uniformly distributed but rather clustered around zero, V = 7.179, p = 0.032 (Figure 4b). Combined with the findings described above, this alignment suggests that, at a given timepoint, the motor activity that encodes temporal predictions can be described as a function of content-specific activity in visual areas (Figure 4c).

## Discussion

Predicting temporal properties of our environment is crucial for our adaptive interaction with it. Here, we show that the human brain represents the temporal structure of the environment in two distinct but complementary ways. We reveal rate-specific phase-coupling of motor areas to the timing of rhythmic events that persists during a period in which events are absent. Oscillations in the absence of external rhythmic stimulation are consistent with entrainment mechanisms that instantiate predictions of event timing and predict the fidelity of subsequent temporal judgements. These temporal predictions are simultaneously embedded in predictions of sensory content; we show that activity encoding stimulus content in visual regions rises and falls in line with temporal predictions. Interestingly, this neural signature of temporally-precise content predictions appears regardless of whether one is explicitly judging time, yet correlates with the strength of motor entrainment when doing so and is phase-aligned.

Our finding of low-frequency ‘entrainment’ in motor regions to visual structure is in line with recent similar findings in the auditory domain, which have led to proposals that temporal prediction is instantiated as simulated action (13–19). The supplementary motor area (SMA), in particular, is proposed to be central to this covert maintenance of rhythms; consistent with the localization of entrained oscillations in our study (Figure 1f). While evidence for analogous processes in human vision had previously been lacking, our findings therefore justify an extension of more-established auditory theories to the visual domain, with a number of important functional implications (20).

Importantly however, our results also contradict some of the central claims of these auditory theories – particularly that sensory predictions are the consequence of (simulated or covert) actions, for example, through efference copies (13,16). Our study showed temporally-precise predictions of sensory content in both time- and content-judgement tasks but that motor-based entrainment was only apparent when time-judgements were required. Therefore, we propose that temporally-precise sensory predictions are unlikely to be the result of motor functioning since we show that they are reliably present even when motor entrainment is absent. Our findings thus lead us to propose an alternative account of how we may represent time. That is, we propose the causally-inverse interpretation – that the visual system continuously represents temporally-precise predictions of future sensory states independently of motor functions, and that the moment-by-moment strength of these visual representations is read out by the motor system if we must directly judge temporal information (Figure 4c, see (21,22)). We thus propose explicit derivation of time to selectively modulate the effective connectivity between visual and motor areas (22,23). This mechanism is more inline with our findings than the existing accounts and could be further tested by examining these processes when motor processing is aberrant, for example in lesion patients (24). More generally, it supports a highly integrated rather than dichotomized neural representation of content and time (25).

Some extant literature is already in line with this interpretation. For example, recent work in non-human primates showed that location-specific firing rates in visual cortex maintain visual rhythms (24, also see 25), and in mice, temporal expectations modulate the activity of neurons tuned to the same frequency as the predicted tone (28). In humans, behavioural and M/EEG work has shown effects of temporal predictions to be reduced in the absence of concurrent sensory (e.g. spatial) predictions (29–33). Further, recent high-field fMRI work showed greater tuning of neural responses to duration and rate at higher levels of the visual processing hierarchy (32, also see 33). On top of said hierarchy, we suggest that motor regions maintain the overall predicted timecourse of information as a model of external temporal structure (Figure 4c). The motor system’s flexibility to represent both periodic and aperiodic temporal structure additionally predicts that such a mechanism would not be limited to isochronous rhythms like the ones used here (19,36).

Our interpretation of temporal predictions as being derived from content-based sensory predictions does not conflict with the idea that the motor system influences sensory processing. This has been the focus of the literature on auditory temporal tracking which has suggested that the motor system’s propensity to synchronize overt and covert action to sensory events leads to improved perception (16, also 35). Our finding that motor-based temporal prediction supports performance in visual timing judgements is consistent with these effects. Sensory predictions supporting motor-based temporal predictions which in turn shape sensory processing would suggest an action-perception-loop. In line with this idea, active sensing theories outline how movement not only passively reflects temporal structure but also reorients sensors to sample the environment in a maximally informative way (38,39).

Finally, we believe that our findings may help to inform controversies concerning the phenomenon of entrainment. While entrained oscillations are a potentially powerful mechanism for tracking and predicting temporal structure, this suggestion has been challenged by scepticism concerning the degree of empirical support (2,3,36). Resolving these inconsistent findings, our study suggests some form of content-specific entrainment whereby, instead of fluctuations in overall neural amplitude, representations of predicted sensory content fluctuate at the rate of external rhythms. If so, the absence of entrainment in measurements that average over the activity of neurons tuned to predicted and non-predicted sensory features would not be surprising. Especially unsurprising in fact, given some research that suggests activating one representation may suppress others, perhaps via lateral inhibition (12,40), thereby not necessarily increasing net activity over time. Interestingly, despite similar suggestions in the past that auditory entrainment may be tonotopically-specific (41,42), neural investigations of entrainment are yet to interrogate this possibility more closely - likely due to difficulties in separating activity from differently-tuned neuronal populations in non-invasive human electrophysiology. Our technique of probing for rhythm-locked content-specific activity may serve as a starting point to overcome these difficulties.

In conclusion, temporal predictions shape our perception of the sensory world. Here, we show that they are neurally represented in two different, but related, ways in sensorimotor hierarchies. In early visual regions, temporal predictions are encoded in continuous temporally-precise predictions of upcoming sensory content: the stronger the representation of the predicted content, the closer a given timepoint matches the temporal prediction. On the other hand, motor regions maintain a representation of temporal predictions when judging temporal features, aligning in phase and magnitude to the content-specific, visual signals. These findings suggest a new way in which sensory and motor encoding may interact to allow us to perceive the intricate temporal relationships in our environment.

## MATERIALS AND METHODS

### Participants

Thirty-one participants were screened for MEG eligibility and gave informed consent in line with the UCL ethics committee (protocol 3090/004). Participants were recruited through the local SONA subject pool, reimbursed in either course credit or cash (£10/hour) and required to have normal or corrected-to-normal vision. One participant was excluded from analyses as they did not complete the experiment, leaving a sample of 30 participants (24 females; mean age = 24.81, SD = 4.39 years; 30 being our preregistered sample size: https://osf.io/8ptsz).

### Stimuli and Task

Gabor patches were created using the Psychophysics Toolbox implemented in MATLAB, with 12° of visual angle diameter and a spatial frequency of 0.4 per visual angle (43,44). Annuli were created by overlaying a background-coloured (grey) circle on its centre that spanned 0.43° of visual angle and contained a black fixation cross of 0.35° of visual angle. Each Gabor was presented for 200ms and the first in the sequence started at one of the 12 possible orientations that result from equally dividing the 0-165° space using steps of 15°.

On 78% of trials within each block, a probe stimulus appeared in time with the preceding rhythm: exactly four frequency cycles from the onset of the final entrainer of the flashing series. On the remaining 22% of trials, it appeared either half a frequency cycle too early or too late (counterbalanced across trials within each block). Furthermore, and independently of this timing manipulation, on 78% of trials the orientation of the probe was the result of adding the fixed amount of rotation from the preceding series to its final patch. On the other 22% of trials, the probe showed either one rotation step added to or subtracted from this typical orientation. Before each of the blocks, participants were told to either judge timing or orientation through text on the screen. In the former, they were asked to press a button when the probe did not appear in time with the preceding rhythm. In the latter, they were asked to press the button when the probe did not show the correct (next following) orientation. They could respond as soon as the probe appeared and had 1500 ms to do so. The next trial started approximately 1250 ms (SD = 300 ms) after the end of the previous trial’s response window. There were three blocks of each stimulation rate and task combination, with presentation order approximately balanced across participants.

All combinations of levels of the other trial variables – number of entrainers, step size, direction and starting angle of orientation rotation as well as direction of deviant probes (forward or backward in rotation or time) – were balanced across trials within a participant and presented in a randomised order.

The study consisted of two testing sessions. The first was a behavioural practice session of one hour in which participants learned and practised both tasks. Given sufficient performance in both tasks as well as MEG eligibility, participants were invited for a scanning session on a different day. If, due to scheduling constraints, the scan occurred more than a week after behavioural practice, participants practised both tasks again for a combined half hour before entering the scanner. In the scanner, participants completed three blocks of 36 trials for each rate and task combination, resulting in 108 trials per condition. We instructed participants to stay still, citing MEG data quality issues as the reason, but also to prevent overt movement along with the stimuli. Participants were monitored during scanning using live video transmission. Following the scan, participants completed a finger tapping task designed to measure their natural motor rates. This measure was recorded as pilot information for a future study and was not analysed for the current study.

### MEG acquisition and preprocessing

MEG data acquisition occurred in a magnetically shieled room using a 272-channel CTF MEG system with axial gradiometers sampling at 600Hz. A third-order gradient compensation algorithm was applied online for noise reduction. Fiducial coils on the nasion as well as on the left and right preauricular were used to monitor the participant’s head position during data acquisition. A photodiode at the bottom left of the screen was used to temporally align the neural signal with on-screen stimulus presentation. Alongside MEG acquisition, an EyeLink 1000 infrared tracker (SR Research Ltd.) was used to record eye movements.

We further preprocessed the MEG data using the Fieldtrip toolbox implemented in MATLAB (45). Visual artifact rejection was carried out using trial-wise MEG sensor variance as the selection criterion for further manual inspection. Trials were removed that showed excessive artifacts. The eye-tracking signal was further used to flag trials in which blinks occurred during stimulus presentation. The resulting data were temporarily downsampled to 200Hz, and a 1-40Hz bandpass-filter (two-pass Butterworth) was applied, before conducting independent components analysis (46). The first 50 components’ topographies and time courses were visually inspected for cardiac and eye-related artifacts, and those components were removed (M = 3.433, SD = 0.219). The resulting projections were applied to the original, not filtered nor downsampled, data.

### Behavioural measures

Behavioural analyses were conducted in MATLAB and focused on the signal detection theoretic (SDT) measure of d-prime (47), calculated as the difference between participants’ normalized hit and false alarm rates within a condition. This measure reflects their sensitivity for detecting the specified target feature in each task.

### Rate-specific intertrial phase consistency

All MEG-related analyses were conducted in MATLAB using the Fieldtrip toolbox. For the following analyses concerning phase consistency, we used custom scripts adapted from (1). We preregistered the analyses mentioned in this section together with the planned sample size, experimental variables and hypothesis on Open Science Framework (https://osf.io/jm6er) before starting data collection. Inspired by past work, we used rate-specific inter-trial phase consistency (ITPC) as a measure of oscillatory tracking. When stimulated at a particular frequency, the neural phase of responses at that frequency should be aligned with the timepoints of stimulus appearance. Since all trials in our design within a specific frequency block follow the same time course of entraining stimuli followed by a maintenance period, we expected that, at a given timepoint, neural activity at the frequency of the entraining stimulus should show a higher ITPC than the alternate frequency. To directly quantify this comparison, we used the rate-specific neural response (RSR) proposed by (1). It quantifies how much more ITPC there is to a given trial’s stimulation rate (e.g. 2Hz) compared to the other, non-stimulated rate (1.3Hz). This approach controls for any non-experimental, spectral differences between the stimulation rates.

To calculate ITPC, preprocessed data were low-pass filtered at 20Hz (two-pass Butterworth) to mitigate the influence of movement-related artifacts. The axial gradiometer data were then subjected to a planar transform, yielding two components for each of the 272 sensors. ITPC calculation was carried out on each of the resulting 544 components separately, after which the two components belonging to each sensor were averaged to restore the original gradiometer structure. We used a sliding-window fast fourier transform (FFT) to transform the data into the frequency domain. The window slid in steps of 20ms with its width set to 1500ms to provide sufficient spectral resolution to distinguish between our stimulation rates 1.33Hz (2 cycles in 1500ms) and 2Hz (3 cycles), and sufficient temporal specificity to isolate responses during the stimulus absent maintenance period. Two cycles providing suitable spectral resolution to reliably detect delta-rate ITPC is supported by recent work – upon which the present study was based – employing a similar analysis and design (1). The phase angle was extracted for both frequencies of interest for each of the timepoints upon which the FFT window was centred.

The complex mean of the resulting trial-by-trial phase was then computed separately over all trials of each of the four (rate by task) conditions before taking the absolute value to yield each condition’s ITPC. Given that ITPC is sensitive to trial count, we confirmed that the four conditions were highly comparable in the number of trials entered into the analysis (M = 94.367, condition-SD = 0.395). To further account for non-oscillatory 1/f-components that exist in the frequency spectrum of neural data and that could bias the RSR, we fitted and subtracted a 1/f curve from the ITPC values for each participant, sensor and condition separately before calculating RSR based on the residuals (1).

Since the number of Gabors in each flashing sequence was the same for both stimulation frequencies – 7 or 8 – the trials in the respective conditions were of different lengths. To align the two, we centred trials of both frequencies to the onset of the final Gabor of the sequence. Then, the longer 1.3Hz trial time courses were cut to the length of the 2Hz trials which allowed us to calculate RSR values for each sensor, task, participant and timepoint upon which the FFT was centred.

To distinguish oscillatory activity from evoked signals, we focused our analyses on timepoints between each sequence’s final Gabor and the onset of the probe. Specifically, we selected the maximal length of data within the three cycles of the stimulus-empty period while avoiding contamination from evoked responses stemming from the offset of stimulation sequence and the onset of the probe. The FFT window centres therefore spanned from 750ms to 1250ms after the onset of the final Gabor. Within this analysis window, RSR values were averaged so that task-wise results from all participants could be statistically evaluated in the following way.

Cluster-based permutation dependent-samples t-tests were used to both compare the two tasks to one another and to evaluate whether the RSR in each individual condition is significantly different (two-sided) from a baseline of 0. Previous work supports this analysis approach by showing that RSR values are normally distributed and that a surrogate distribution based on adding random phase values to the FFT output before calculating ITPC is indeed centred around 0 (1). We used a Monte-Carlo algorithm with 5000 permutations for each planar-transformed MEG sensor separately. Clusters were formed by neighbouring (minimum 2) sensors which individually reached significance (alpha = 0.05). The resulting cluster-level statistic represents the sum of the t-values of the sensors within the cluster and is tested for significance (cluster-alpha = 0.05).

### Source localisation

In order to identify the source of the RSR signal, as well as for visualization purposes, we conducted source localisation by using a template approach as implemented in Fieldtrip. Using participants’ fiducial measurements and the MEG sensor structure, we warped a standard Fieldtrip single shell head model and a standard Fieldtrip source model (8mm dipole spacing) into participant-specific approximations. We estimated a spatial filter for the analysis window described above using LCMV-beamformers after combining the data from the two stimulation rates (48). We applied the spatial filters to FFT-transformed single-trial data before extracting the phase and calculating ITPC and RSR as in the sensor-level data described above. We used the AAL atlas implemented in Fieldtrip for probabilistic anatomical labelling (5).

### Orientation decoding

We used an inverted encoding model (IEM) to decode sensory representations, specifically an implementation used in previous work that was able to decode predicted grating orientations in the absence of ongoing stimulus presentation. We will provide a condensed description of the IEM here but for a full implementation, see (10) and (11), from which we used custom MATLAB scripts.

At its core, the approach includes two steps. First, a forward model is trained to model the MEG sensor-level activity evoked by different Gabor orientations. Specifically, this model consists of 32 equally-spaced channels whose Von Mises tuning curves cover the entire orientation spectrum and can be linearly combined to reproduce any single orientation. Once trained, the transition from orientation to MEG data is described by a weight matrix that captures each MEG sensor’s ‘orientation tuning’ while accounting for the noise covariance of neighbouring sensors. Note that we also examined this weight matrix to probe which sensors contribute to the decoder (Figure 3a). In the second step of the IEM, this filter is applied in the opposite direction to create an inverse model that converts MEG data into evidence for the orientation spectrum.

In this study, we used the first two Gabors of each trial’s flashing sequence to train the forward model. We chose the first two Gabors because their orientation could not be predicted making their neural response the result of ‘bottom-up’ processing only, in line with the logic behind functional localizers (49). After the second Gabor is presented, participants can anticipate subsequent Gabor orientations since the initial orientation, rotation direction and stepsize for that trial have already been determined. Decoding did not improve when adding more than the first two Gabors of each sequence to the training data. We extracted two 500ms epochs from each preprocessed trial, starting at the respective onsets of the first and second Gabor of the sequence. We also applied a 40Hz low-pass filter with a filter order of 6, in line with past research (10). This process yielded an average of around 840 (SD = 34) training examples (∼ 70 per orientation class) per participant. The model was trained at each training data timepoint, separated by 5ms. While doing so, the data were averaged within windows of 29.2ms using a sliding window approach to improve the signal-noise-ratio at each training timepoint (10). The resulting time-resolved filter matrix was then applied to our test data.

We again focused our analysis on the time window between the final entrainer and the probe onset. Specifically, we extracted the epoch between each trial sequence’s final Gabor and 1000ms after the probe onset and again applied a 40Hz low-pass filter with a filter order of 6. We estimated the orientation channel responses of these test data at each timepoint, separated by 5ms and averaged within a window of 30ms (30ms minus one sample to assure asymmetry, in line with (10)) - which yielded a 4D matrix (training time X testing time X model channel X trial) containing the estimated channel responses during the window of interest for each trial. Channel responses were consequently shifted so that the channel centred on the orientation of interest (see below) occupied the position of the 0°-centred channel. This allowed averaging over trials to result in a decoding output of the orientation of interest which was consequently transformed into a scalar projection that was used as the measure of decoding performance. Trial-averaging was done separately for each condition (orientation/timing task) and rate (1.3Hz/2Hz).

We determined four orientations of interest for every trial. Namely, based on each trial’s orientation sequence parameters (starting orientation, rotation direction and rotation step size), we determined the three orientations that would have appeared in the three cycles of the stimulus-empty window if the sequence had continued. These could subsequently be used to establish whether neural activity represented the predicted continuation of the previous stimulus sequence. The fourth orientation of interest was the presented probe orientation, which was used to establish the fidelity of the decoding approach. We focused decoding analyses exclusively on expected trials since unexpected presentations or omissions could infringe on the analysis window or lead to divergent responses. This resulted in testing data based on an average of around 66 trials per participant per condition (rate by task).

### Content-specific temporal prediction analysis

Within the decoding output, we selected a training time window from 120 to 160ms after stimulus onset, based on past research showing optimal decoding sensitivity for similar stimuli and presentation durations in this window, and averaged the decoding output within this window (10). We first tested the approach’s fidelity using the probes presented in both tasks to establish that orientation signals can be decoded at above-chance performance. Then, the decoding output of each trial was narrowed to the extrapolated orientations during the time between the onsets of the final sequence Gabor and the probe. This resulted in windows of 2000 or 3000ms for decoding output with 2 or 1.3Hz stimulation rate respectively. This yielded three decoding traces for each trial, one for each extrapolated orientation. To focus on delta-rate representation changes and to improve the signal-to-noise ratio, we next smoothed these traces by averaging with a 150ms sliding window (50). Finally, we interpolated the decoding output from our two stimulus presentation rates onto a common time axis of rate-specific radians.

We next defined a linear mixed effects model in which decoding evidence of the three extrapolated orientations was predicted from content predictions (the orientation that would have appeared in the timepoint’s frequency-specific cycle), temporal predictions which were modelled as a cosine that peaks at the respective onsets of the predicted Gabors, task as well as all interactions between these predictors.

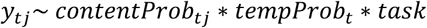

where *y* is the decoding evidence for orientation *j* at timepoint *t*. *contentProb* is a binary vector (expected / unexpected) that denotes for each orientation *j* whether it is the most likely orientation at timepoint *t*. *tempProb* is the zero-lag cosine transformation of time in radians. *task* is a binary vector (orientation / timing). Random intercepts were added for each of the three possible orientations along with random slopes and intercepts for each participant.

We fitted this model to each participant, separately for each task using the lme4 package in R and used the afex package to retrieve p-values for predictor slopes (51,52). Marginal means were estimated for the two levels of the content prediction term as well as the extreme terms of the temporal prediction (i.e. -1 and 1) using the emmeans package in R (53). For illustration purposes, the empirical group-averages displayed in Figure 3b were z-scored against each decoding output’s mean and standard deviation. To quantify the temporal precision of content predictions that was correlated with each participants’ RSR, we first computed the content prediction effect as the difference in marginal means between the predicted and unpredicted content at the peak (cos(t) = 1, hence t = 2π, 4π, 6π, in Figure 2a) and at the trough (cos(t) = -1, 3π, 5π, etc) of the temporal expectation respectively. Then, the temporal precision of content predictions was quantified as the difference between the content prediction effects at the peak compared to the troughs. This equates to the difference between the two columns in each task in Figure 3d. Note that all statistical patterns were replicated if defining the content prediction effect simply according to predicted orientations or interaction beta values.

### Orientation decoding source localisation

To determine the neural sources of the IEM-based decoding of Gabor orientations, we focused on the weight matrix of the forward model that maps MEG sensor activity onto the model’s hypothetical orientation channels for each training timepoint (see above). We used LCMV beamforming as we did for the ITPC analyses, but this time created the spatial filter based on the epochs that served as training data for the IEM. We multiplied each training timepoint’s IEM weight matrix with the resulting source filter before taking the absolute value of the product and averaging over the model’s hypothetical orientation channels. Finally, we averaged each voxel’s model weight within our IEM training window of interest. To account for the centre of head bias that arises during source localisation, we repeated the above steps using a sensor-wise shuffled version of the IEM weight matrix so that the source filter was multiplied with a permuted weight matrix. Specifically, we ran 100 repetitions of this procedure, averaged the result and consequently z-scored the true weight matrix against this shuffled baseline. Since the permuted baseline still contains the centre of head bias after multiplication with the spatial filter but no longer any meaningful weights, this procedure results in a bias-free source-localisation of the IEM weight matrix.

### Visuomotor coupling analysis

In order to assess the coupling between rate-specific motor phase and content-specific visual activity, we first had to quantify the temporal profile. We did so using a cosine-lagged regression approach as described below. The rate-specific motor phase represented the sensor and trial-averaged phase within the significant timing task RSR cluster (Figure 1c) for the stimulation-congruent frequency (e.g. 2Hz phase for 2Hz stimulation rate) during the stimulus-empty window (-0.25s before final sequence Gabor onset to 0.25s after probe onset). Before analysis, phase values were subjected to a cosine-transform as displayed in Figure 1e. The content-specific visual activity concerned the content prediction effect (predicted – unpredicted) as displayed in Figure 3d that we had previously modelled as a fixed, zero-lag cosine coupled to the predicted onsets during the stimulus-empty window (Figure 2).

In this analysis, we varied this cosine lag to test which temporal lags best described this content prediction effect. Separately for each stimulation rate, task and participant, we predicted the motor and visual signals from a family of 20 cosines (steps of pi/10) that were shifted up to half a cycle (1.33Hz: 750ms, 2Hz: 500ms) backwards or forwards in time relative to the predicted onsets. For the motor signal, this was done through a simple linear model where the cosine-transformed phase was predicted from a rate-specific, iteratively-shifted cosine and the standardized beta coefficient was recorded. For the visual signal, we simply replaced the temporal prediction term of the mixed model (Figure 2, here fitted separately for each participant and task for computational reasons) with the rate-specific, iteratively-shifted cosine, refit the model and calculated the temporal specificity of content predictions like before (peak – trough, Figure 4d) to capture how well the given cosine described the content prediction effect in visual activity. After averaging over the two stimulation rates, the effects of interest across the 20 tested cosines yielded lag-effect time courses describing the rate-specific motor phase and the content-specific visual activity respectively. These time courses were statistically evaluated using cluster-based permutation tests and visually displayed in the supplemental materials (Figure S4).

To test the coupling between these two signals within each participant, we used lagged cross-correlation as implemented in the ‘acf’ function of the R package ‘stats’ (54). We did so only for the timing task where both motor and visual signatures were present (also see Figure S4). For each participant, we retrieved the cross-correlation lag that yielded the highest correlation between the two signal time courses. Negative lags represented the visual leading the motor signal and positive lags represented the motor leading the visual signal. We tested for uniformity in visuomotor lags within our sample using a circular V test with the alternative hypothesis of clustering around 0 (55).

## Acknowledgments

The authors would like to thank the FIL’s imaging support team, particularly Dimitra Moraiti and Daniel Bates, as well as Dorottya Hetenyi for MEG scanning assistance. This work was supported by a Leverhulme Trust project grant (RPG-2022-358) and European Research Council (ERC) consolidator grant (101001592) under the European Union’s Horizon 2020 research and innovation programme, both awarded to CP, and Medical Research Council funding of the Cognition and Brain Sciences Unit (MHD, MC_UU_00030/6). The Department of Imaging Neuroscience was supported by core funding from the Wellcome Trust (203147/Z/16/Z).

## Author contributions

Conceptualization, A.K., Q.G., P.K., M.H.D, C.P.; data curation, A.K.; formal analysis, A.K.; funding acquisition, C.P.; investigation, A.K.; methodology, A.K., Q.G., P.K., M.H.D, C.P.; project administration, A.K., C.P.; software, A.K.; resources, C.P.; validation, A.K.; visualization, A.K.; writing – original draft, A.K.; writing – review and editing, A.K., Q.G., P.K., M.H.D, C.P.

## Notes

### Competing Interest Statement

The authors have declared no competing interest.

### Summary of Updates

Additional analyses relating to motor and visual signatures to one another as well as further control analyses.

