## Supplemental materials for "Temporal predictions as motor readouts of sensory predictions"

**Behavioural data**

We present in Table S1 all d-primes for all cells of our design, along overall accuracy, hit and false alarm rates as well as RTs. We also compared the d-primes for early vs. late violations in the timing task, separately for the 1.33Hz and 2Hz conditions. We did not find a significant difference in the 1.33Hz, $t(29) = -1.377$, $p = 0.179$, nor in the 2Hz condition, $t(29) = 0.650$, $p = 0.521$. The opposite trends between the two rates resulted in no difference when collapsing over rates, $t(29) = -0.451$, $p = 0.656$. We did not find a response time (RT) difference between the two tasks, paired t-test of RTs: $t(29) = -1.331$, $p = 0.194$.

The most important element of the task performance, given our question, was that participants were effectively performing the tasks above chance. However, given the difference in task performance between task and rate, we double checked that it was not responsible for task differences in neural effects. We first investigated whether higher difficulty in the timing compared to the orientation task may be responsible for the relatively larger RSR. We deemed this possibility low, given the significant correlation between timing task performance and RSR ($r = 0.385$, $p = 0.036$, Figure 1g): participants with higher timing task performance show the strongest RSR.

Next, we considered the converse - whether the RSR did not emerge in the orientation task because it was easier. Rate-specific (1.33 and 2Hz) d-primes of 1.516 and 1.348 in the orientation task are do not indicate ceiling effects, but we nevertheless the tested possibility by creating a subset of 10 participants with the lowest orientation task d-primes ($M = 0.784$, $SEM = 0.113$). If high task difficulty generated an RSR, we should find evidence for an orientation task RSR in this subsample. We therefore reran our cluster-based permutation test on this subsample. We did not find any candidate clusters for RSR in the orientation task in this subsample (Figure S1a, left). This result can not be explained by the low power in this sample of 10 participants since we were able to recover the timing task RSR effect, cluster-corrected $p = 0.049$; summed $t = 26.827$, 9 sensors (Figure S1a, right). Difficulty alone is thus unlikely to explain the difference in RSR between the two tasks.

Third, we probed whether participants who exhibited poor performance in the timing task did not meaningfully attempt the task, thereby explaining the correlation between RSR and timing task performance. Contrary to this notion, even the 10 poorest timing task performers ($M = 0.032$) still showed a significant RSR within the cluster of sensors that emerged from the whole sample analysis, t-test of participant timing task RSRs against 0: $M = 0.026$, $t(9) = 2.500$, $p = 0.034$. Note that this is despite the positive relationship between RSR and performance. It is therefore unlikely that the link between timing task RSR and behavioural performance could be explained by a subset who were disengaged.

To further test this possibility, we tested for the relationship between RSR and behavioural performance in the 86 (matching number of sensors in timing task cluster of the stimulus-empty window) sensors that showed the strongest RSR response during stimulation (sequence viewing). Note that RSR in this time window is likely attributable to rhythm-locked evoked responses, consistent with the strongest-responding sensors being located in occipital areas (Figure S2a). General inattentiveness, or in extreme terms, participants closing their eyes, should lead to a reduced response in this window's RSR as past research has shown rate-specific phase-locking in sensory areas to depend on attention (1). We were unable to find a relationship between RSR during stimulation and behavioural performance in the orientation task ($r = -0.229$, $p = 0.224$, Figure S1b), nor in the timing task ($r = -0.156$, $p = 0.410$, Figure S1b). RSR strength, and therefore the timing task relationship between maintained RSR and behavioural performance, is therefore unlikely to be explained by differences in general attentiveness.

As outlined above, we see the most crucial element of the behavioural data the demonstration that participants could perform the two tasks above chance, but these further analyses also provide assurance concerning comparisons across tasks.

| | 1.33Hz Orientation | 1.33Hz Timing | 2Hz Orientation | 2Hz Timing |
|---|---|---|---|---|
| D-Prime<br>Mean (SEM) | 1.516 (0.126) | 0.667 (0.111) | 1.348 (0.102) | 0.425 (0.099) |
| Accuracy<br>Mean (SEM) | 0.816 (0.017) | 0.737 (0.013) | 0.807 (0.016) | 0.732 (0.010) |
| Hit rate<br>Mean (SEM) | 0.575 (0.033) | 0.363 (0.030) | 0.497 (0.025) | 0.259 (0.029) |
| False alarm rate<br>Mean (SEM) | 0.117 (0.020) | 0.157 (0.014) | 0.106 (0.021) | 0.136 (0.012) |
| RT (s)<br>Mean (SEM) | 0.858 (0.024) | 0.892 (0.044) | 0.819 (0.027) | 0.871 (0.041) |

**Table S1 Raw behavioural measures across all task and rate conditions.**

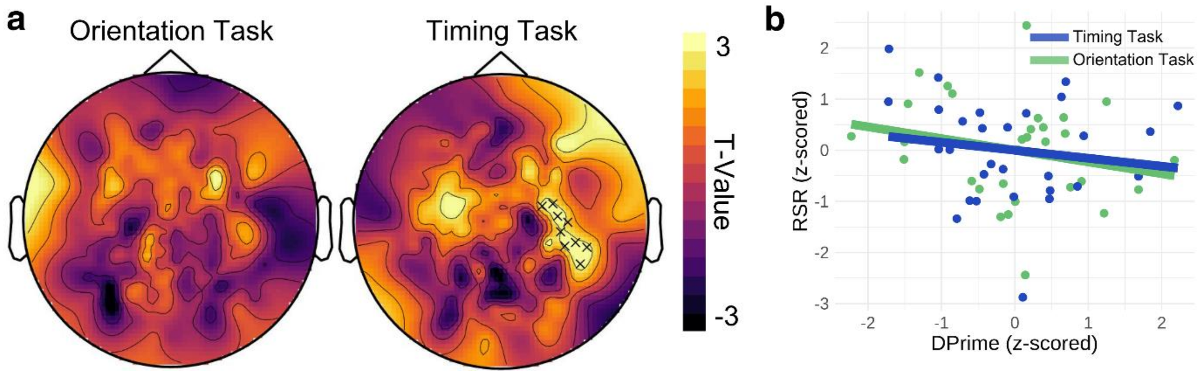


**Figure S1 Additional behavioural analyses controlling for task difficulty. a** Permutation test t-value topographies for the RSR during the Maintenance window, separately for the two tasks in the subset of participant with lower orientation task performance. Sensors marked with a cross belonged to the significant cluster found for a given condition. **b** Correlations between rate-specific neural activity averaged across the sensors showing the strongest RSR during sequence (see S2) and task performance, separately for the orientation and timing task. Each circle represents a single participant (N = 30) in the given task.

### RSR during stimulation

The RSR is presented below for a time window during ongoing visual sequence presentation. Specifically, we considered the time window from 2.25s before to the onset of the final sequence Gabor. In line with an interpretation of repeated visual evoked responses locked to each sequence Gabor, we found significant RSR in both tasks (orientation task: cluster-corrected $p < 0.001$; summed $t = 3086.2$; 272 sensors, timing task: cluster-corrected $p < 0.001$; summed $t = 3118.0$; 272 sensors) with strongest effects in sensors over occipital areas but no reliable difference in RSR between the two tasks (all cluster-corrected p > 0.05, Figure S2a).

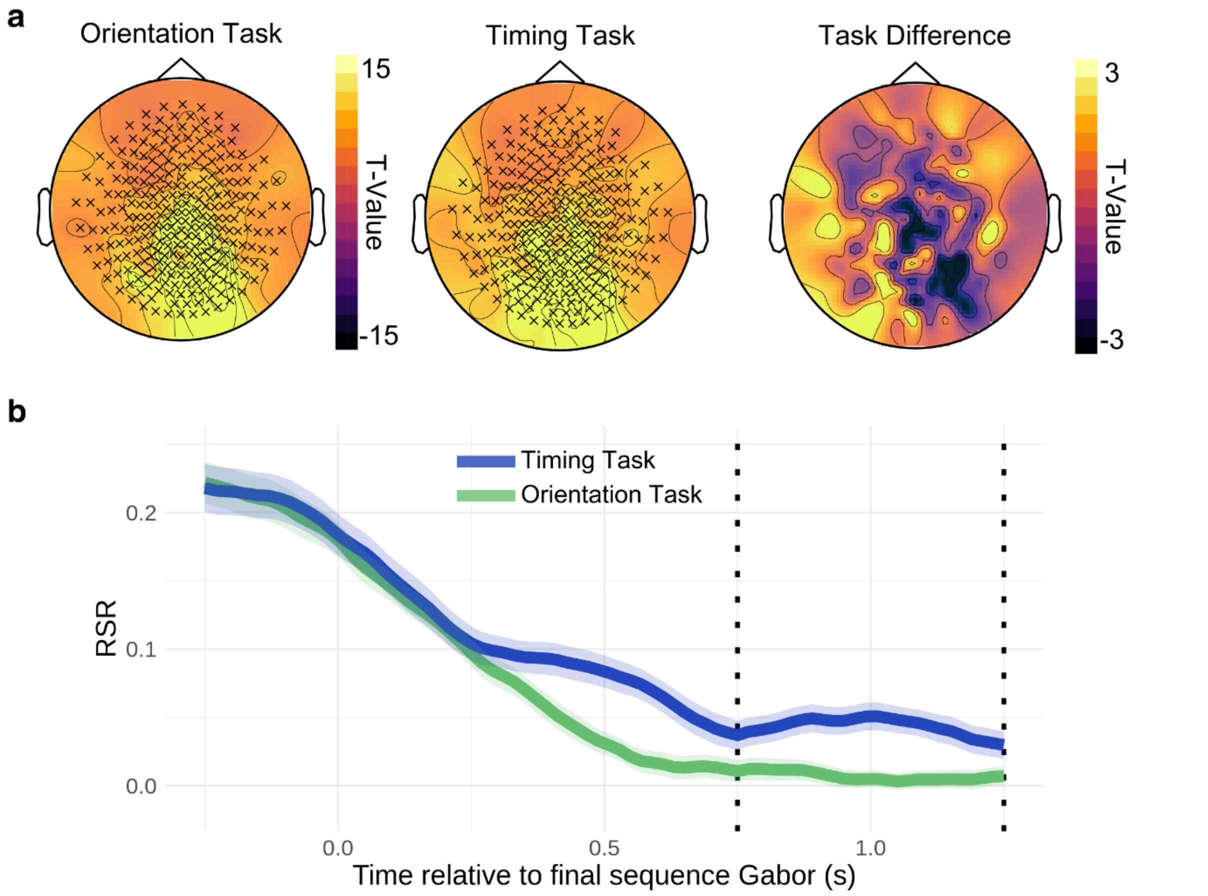


**Figure S2 a** Permutation test t-value topographies for the RSR during ongoing visual stimulation, separately for the two tasks and for the task difference. Sensors marked with a cross belonged to the significant cluster found for a given condition. **b** Time course of RSR in significant motor sensors (Figure 1c) from end of sequence into the stimulus-empty window. Dotted vertical lines signal the beginning and end of the designated Maintenance window upon which the main RSR analysis focused. Each task's line represents the group mean and the surrounding shaded area its standard error.

#### Testing for overt responses

We aimed to minimize simultaneous tracking movements by monitoring participants through live video transmission during the scan and by emphasizing before every block that movement would render the recorded neural data unusable. However, we also investigated the possibility of simultaneous movement using our neural data. To this end, we conducted an evoked response analysis within the sensors that showed the significant RSR in the timing task. This is based upon past research showing that overtly moving along to sensory rhythms, but not passively tracking them, leads to rhythm-locked event-related fields (ERFs) in the MEG signal (2). Separately for each participant, task and stimulation rate, we time-locked activity to the onset of the final sequence Gabor, averaged over trials and computed the planar-transform. To isolate rhythm-locked ERFs in our frequencies of interest, we removed slow drift signals in the output by high-pass filtering at 2/3 Hz. We consequently used cluster-based permutation testing to test for rhythm-locked activity in individual conditions and in the difference between tasks.

Cluster-based permutation testing revealed no rhythm-locked, motor-evoked activity in the stimulus-empty window in any rate or task condition. Within the considered time window, significant activity only appeared in the visually-evoked response to the final sequence Gabor, not in the stimulus-empty window that we analysed for the RSR.

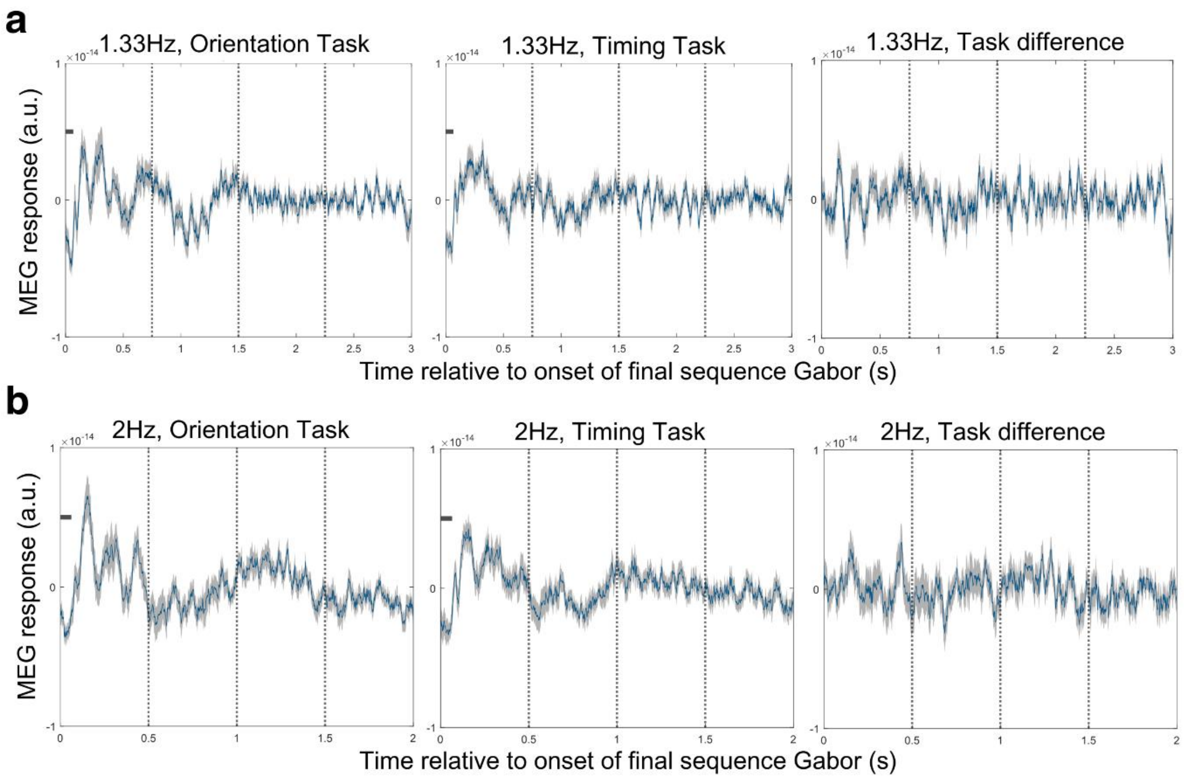


**Figure S3** Motor-evoked event-related fields (ERFs) averaged over sensors belong to timing task RSR cluster (Figure 1c) for **a** 1.33Hz conditions and **b** 2Hz conditions, each displayed separately for orientation task (left), timing task (middle) and task difference (right). Horizontal black bars reflect statistically-significant clusters which can only be found in the visually-evoked response to the final sequence Gabor, does not differ between rates and is therefore unlikely to explain the motor RSR. Blue lines represent the group mean and the surrounding shaded area its standard error.

A more indirect indicator for the lack of overt movements in our timing task is derived from recent work by Zalta, Petkoski and Morillon (3). The authors investigate within which frequency ranges overt movement aids the tracking of sensory rhythms in the auditory and, more importantly for us, visual domains. They found that within the delta range that we investigated (1.33 to 2Hz), participants' temporal deviation task performance was worse when they overtly tapped along to the visual rhythm compared to passively tracking it. The authors concluded that overt motor contribution disrupts visual temporal performance in these tempi. If our RSR response reflected overt movement, the strength of this signal may therefore be expected to correlate negatively with timing task performance. Given that it does not, and instead correlates positively, is more in line with a passive tracking of visual rhythms.

Finally, we suggest that popular accounts positing passive rhythmic tracking as action simulation would deem the distinction between overt movement and passive tracking a quantitative, rather than qualitative one. The idea is that both processes engage similar neural patterns and whether a movement is truly carried out or not is a matter of passing a quantitative threshold that elicits overt effector movement or not. Our proposal of motor temporal predictions representing readouts of sensory predictions builds on these accounts by assuming that the motor temporal prediction can take either overt or covert form.

**Lagged regression analyses**

The lagged regression of motor phase and content-specific visual activity resulted in lag-effect time courses (see black line in S4, also see Methods). Cluster-based permutation testing over all tested shifts ($-\pi$ to $\pi$ in steps of $\pi/10$) was used to identify lags at which the cosine predictor significantly explained each signal. For the motor phase, we found a significant positive cluster from 1.561 to 0.314 radians before expected stimulus onsets in the timing task (cluster-corrected $p = 0.012$, Figure S4a right). The maximal beta was found for the cosine peaking 0.942 radians before the predicted onsets (vertical lines in Figure S4). This corresponds to 0.113 and 0.075 seconds for the 1.33 and 2Hz conditions respectively. Motor RSR thus anticipates the expected stimulus onsets which is evident in S4a where strongest effect lags peak just before the vertical lines representing the expected onsets. No cluster was found for the orientation task, in line with the lack of an RSR in this task (Figure S4a left).

For content-specific visual activity, we found significant positive clusters in both tasks. In the orientation task, the cluster spanned lags from 0.628 radians before to 1.257 radians after expected stimulus onsets (cluster-corrected $p = 0.036$, Figure S4b left). In the timing task, the cluster similarly spanned from 1.257 radians before to 1.257 radians after expected stimulus onsets (cluster-corrected $p = 0.012$, Figure S4b left). Radians of 0.628 and 1.257 correspond to 0.075 and 0.150 seconds in the 1.33Hz condition and 0.050 and 0.100 seconds in the 2Hz condition. Note that the effect at the cosine lag of zero corresponds to the effect described in the Main Text since we assumed a zero-lag in our original model.

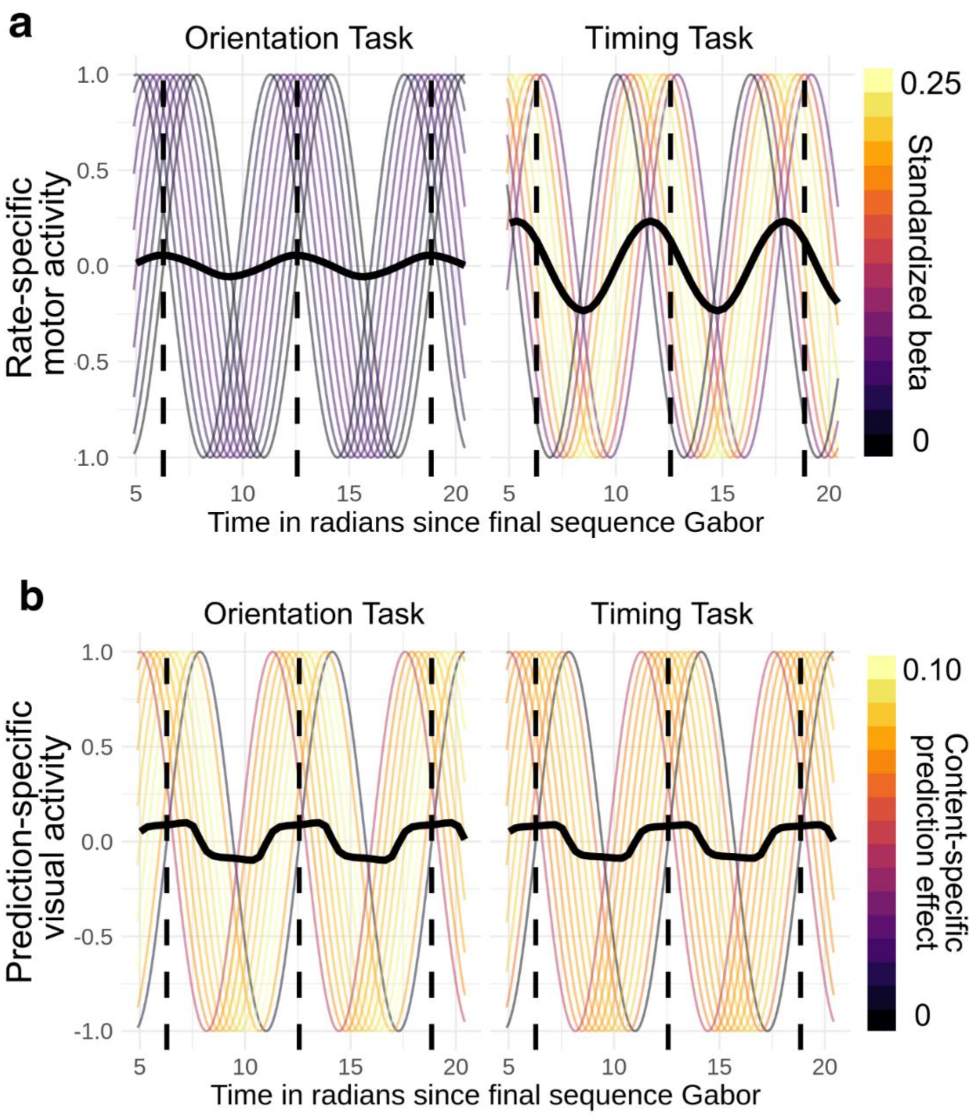


**Figure S4** Lagged regression results for **a** Rate-specific motor phase and **b** content prediction- specific (predicted – unpredicted) visual activity. Thick, black lines represents effect of interest at each tested lag, copied over cycles for illustration purposes. For **a** and **b**, lags that produced negative effects are omitted from visualization since they are equivalent to the additive inverse of the displayed positive-beta cosines shifted by half a cycle, due to cosine periodicity.
